# Regenerated Interneurons Integrate Into Locomotor Circuitry Following Spinal Cord Injury

**DOI:** 10.1101/2020.03.23.003806

**Authors:** Deeptha Vasudevan, Yen-Chyi Liu, Joshua P. Barrios, Maya K. Wheeler, Adam D. Douglass, Richard I. Dorsky

## Abstract

Whereas humans and other adult mammals lack the ability to regain locomotor function after spinal cord injury, zebrafish are able to recover swimming behavior even after complete spinal cord transection. We have previously shown that zebrafish larvae regenerate lost neurons within 9 days post-injury (dpi), but the functional contribution of these neurons to motor recovery is unknown. Here we show that multiple interneuron subtypes known to play a role in locomotor circuitry are regenerated in injured spinal cord segments during the period of functional recovery. Further, we show that one subtype of newly-generated interneurons receives excitatory input and fires synchronously with motor output by 9 dpi. Taken together, our data show that regenerative neurogenesis in the zebrafish spinal cord produces interneurons with the physiological capacity to participate in the recovery of locomotor function.

## Introduction

Spinal cord injury (SCI) is a debilitating condition that leads to loss of motor function due to the severing of axonal connections and the death of neurons at the injury site (Darian-Smith 2009, Jutzeler et al 2019). While functional loss is largely permanent in adult mammals due to an inability to repair damaged tissue, teleost fish and urodele amphibians are able to recover normal locomotor behavior even after complete spinal cord transection (Becker et al 1997, Sirbulescu & Zupanc 2010). In zebrafish, recovery coincides with regrowth of ascending and descending axons from neurons distal to the injury, as well as the regeneration of neurons at the injury site (Becker et al 1997, Briona & Dorsky 2014). Despite evidence for a potential role of local neurogenesis in functional recovery (Duan et al 2016, Jorstad et al 2017, Kumamaru et al 2018), strategies to restore motor function in human SCI patients and animal models have focused primarily on axon regrowth (Anderson et al 2018, Magnusson et al 2014, Mokalled et al 2016). Determining whether newly-generated neurons can integrate into functional circuitry after SCI is therefore important for understanding the regenerative potential of the central nervous system, as well as for designing future therapeutic approaches.

Larval zebrafish injured by surgical spinal cord transection at 5 days post fertilization (dpf) show axon regrowth by 3 days post injury (dpi), neurogenesis by 5 dpi, and recovery of swimming behavior by 9 dpi (Briona & Dorsky 2014). Resident radial glial cells proliferate and migrate longitudinally into the injury site following SCI, and form a glial bridge that acts as a scaffold for ascending and descending axons to regrow (Goldshmit et al 2012). Previous work from our laboratory has shown that these same radial glia also undergo differentiation into neurons that populate the injury site (Briona et al 2015). However, the specific role of radial glia-derived neurons in functional recovery is unknown.

Functional recovery of forward swimming behavior in larval zebrafish requires the re-establishment of rhythmic left-right alternations of trunk axial muscle contractions, which propagate in a rostral-caudal wave (Bagnall & McLean 2014, Gabriel et al 2011). As in other vertebrates, alternating rhythmic motor activity is controlled by a central pattern generator comprising local spinal interneurons (Goulding 2009). The loss of these interneurons and their connections in SCI thus disrupts both rostral-caudal propagation and left-right alternation at and below the injured segment. While regrowth of ascending and descending axons across the injury site may partially restore motor activity in caudal segments, it is also possible that regeneration of local interneurons may participate in the restoration of wave propagation for efficient forward movement.

In this study we used genetic, electrophysiological, and behavioral approaches to examine how the identity and physiology of neurons generated after SCI relates to functional recovery. We found that neurogenesis in transected spinal cord segments correlates with partial recovery of motor wave propagation and left-right alternation by 9 dpi. Furthermore, our data show that newly-generated spinal interneurons comprise identified subtypes known to participate in motor circuits, and that one subtype of these interneurons receives synaptic inputs and fires synchronously with evoked motor output. Together, our results demonstrate for the first time that regenerative neurogenesis produces interneurons with the physiological capacity to participate in spinal cord function after injury.

## Materials and Methods

### Use of Zebrafish

The following zebrafish strains were used in this study: *AB, *mitfa*^*w2*^ (Lister et al 1999), *Tg(elavl3:GAL4)*^*zf409Tg*^ (Chen et al 2012) *Tg(UAS:E1B-Kaede)*^*s1999tTg*^ (Scott et al 2007), *Tg(tph2:Gal4FF)*^*y228Tg*^ (Yokogawa et al 2012), *TgBAC(vsx2:GAL4FF)*^*nns18Tg*^ (Kimura et al 2008), *Tg(5xUAS:eGFP)*^*zf82Tg*^ (Asakawa et al 2008), and *Tg(−20*.*7gata2:EGFP*)^*la3Tg*^ (Meng et al 1997). All fish were raised and bred according to standard procedures (Westerfield 2000). All experiments were approved by the University of Utah’s Institutional Animal Care and Use Committee (IACUC). Fish were crossed and staged according to Kimmel et al., (1995).

#### Spinal cord injury

Larvae were anesthetized at 5 dpf in 0.016% Tricaine (Sigma), mounted on a Sylgard dish and injured by complete transection of the spinal cord at the level of the anal pore as described previously (Briona & Dorsky 2014), except that a sharpened steel pin (FST 26007-05) was used for injury, and recovering larvae were fed rotifers starting at 1 dpi.

#### Kaede photoconversion

Larvae were mounted laterally in 1.5% low-melting agarose in E3 medium. Under a compound microscope with a 20X objective, 405nm light was used to irradiate and photoconvert approximately 20 segments in the trunk, centered on the prospective transection site in the segment dorsal to the anal pore. Larvae to be analyzed were mounted in 1.5% low-melting agarose in E3 laterally and imaged on a Zeiss LSM 880 confocal microscope with AiryScan processing. From maximum-intensity z-projections, a 120*µ*m wide box was drawn centered on the injury plane and red and green neurons were counted using Fiji. Identification of red neurons was based on fluorescence intensity above the threshold of unconverted Kaede.

#### Immunohistochemistry

Larvae were fixed in 4% paraformaldehyde overnight at 4°C, then blocked with goat serum for 1.5 hours, followed by overnight primary and secondary antibody incubation. Stained fish were cleared in 80% glycerol overnight before being mounted on slides with Fluoromount-G (Southern Biotech). Primary antibodies: Rabbit anti-GFP (1:5000, Invitrogen #A-11122), chicken anti-GFP (1:1000, Aves #GFP-1020) and rabbit anti-Pax2a (1:200, Genetex #GTX128127). Secondary antibodies: goat anti-rabbit 488 (1:200, Invitrogen #A-11008), goat anti-chicken 488 (1:200, Invitrogen #A-11039).

#### Recording of swim behavior

Larvae at 3 and 9 dpi were head-embedded in 1.5% low-melting agarose in E3, leaving the regions caudal to the swim bladder free to move. Fish were imaged from above at 500 frames per second using a high-speed Pike F-032B/F-032C VGA camera. For the optomotor response assay, moving black and white gradient bars with an interval of 1 cm and speed of 1cm/sec were projected onto the stage beneath the fish (Orger et al 2000). Ten total trials of 10s were recorded per fish, with a 20s interval between trials.

#### Behavioral analysis

Tail bend angle was measured using a series of 20 tracking points, from the end of the swim bladder to the tip of the tail, and the injury site was specified manually. After extracting 59 kinematic parameters from each discrete behavioral bout, a self-organizing map clustering algorithm was used to classify behaviors. Clusters representing forward swims were clearly identifiable as low-amplitude sinusoidal tail bends with a maximum bend angle less than 90 degrees. The power spectral density (PSD) of tail bends during forward swims was then analyzed separately in regions of the tail rostral and caudal to the injury site. Paired T-tests were performed between uninjured and injured fish for rostral and caudal regions at 3 and 9 dpi at the physiologically relevant range of 10-30 Hz, determined from swim behaviors in uninjured larvae.

### Electrophysiological analysis

Zebrafish larvae were immobilized with a 1mg/mL α-bungarotoxin (Thermo Fisher Scientific) solution for 3-5 minutes and rinsed off thoroughly with extracellular solution (134 mmol NaCl, 2.9 mmol KCl, 1.2 mmol MgCl2, 2.1 mmol CaCl2, 10 mmol HEPES, 10 mmol glucose, pH 7.8). Fish were stabilized on Sylgard dishes with tungsten pins and kept in extracellular solution throughout the experiment. For electrical stimulation, bipolar electrodes were embedded in Sylgard and contacted the head of larvae. Brief electrical shocks of 0.08-0.2ms, with an amplitude of 6-14 V, were applied through an external stimulator (Grass) to elicit fictive swimming behavior. Changes in light levels through adjusting illumination levels on the microscope were also used to elicit a wider range of swim frequencies for all experiments. For peripheral motor nerve recordings, extracellular suction electrodes were made from borosilicate glass capillaries with an outer diameter of 1.5 mm and inner diameter of 0.84 mm (World Precision Instruments) using a micropipette puller (HEKA PIP 6). Electrode tips were beveled to 30-45 degrees with an aperture opening of 20-50 *µ*m using a motorized micropipette grinder (Narishige EG-44).

To access neurons in the spinal cord for whole-cell patch recordings, dorsal musculature was removed from the segment of interest and adjacent rostral and caudal segments. Patch electrodes were made using the same glass capillaries and equipment described for extracellular electrodes and were filled with patch solution (125 mmol K-gluconate, 4 mmol MgCl2, 5 mmol EGTA, 10 mmol HEPES, 4 mmol Na2ATP, pH 7.2). 50 *µ*M Alexa Fluor 568 hydrazide (Thermo Fisher Scientific) was included in the patch solution to label neurons. Patch electrodes had resistances between 12-16 MΩ, and positive pressure was maintained at ∼25 mmHg with a pressure transducer (Fluke DPM1B) during approach to prevent blockage of the tip. Electrophysiological recordings were acquired with a MultiClamp 700B (Molecular Devices) and DigiData 1440A (Molecular Devices), controlled with pClamp 10 software (Molecular Devices). Data were analyzed using Clampfit (Molecular Devices) and DataView (Heitler 2009).

To correlate motor activity to neuronal activity patterns in injured fish, motor output was recorded from the same segment as neuron recordings. However, in injured fish, motor output was recorded from segments rostral to the injury in order to obtain motor patterns that had not been compromised by injury. In injured fish, motor patterns at the injury segment were derived from the recorded motor pattern of the anterior segment adjusted by the average propagation delay per segment calculated from uninjured fish (1.48 ± 1.18 ms/segment, n=26 swim bouts). Motor output patterns recorded from rostral segments in the injured fish were then time-shifted according to how many segments separated the recording electrode from the injury segment to obtain the calculated motor output at injury segments. The derived motor output was used for injured fish for all corresponding analyses.

Motor output recordings were analyzed if a swim bout contained more than two swim bursts recorded per electrode following the stimulus. Motor bursts were detected using DataView, and defined by signal above the baseline noise threshold and separated from the following burst by more than 10 ms. Burst durations and onset latencies between bursts were calculated using Dataview. Spikes were also detected by setting a threshold and spike frequency and phase was calculated using DataView. Statistical analyses were performed with Microsoft Excel. Since data were not normally distributed, the Kolmogorov-Smirnov test was used to determine statistical significance.

## Results

### Motor patterns for rhythmic swimming behavior partially recover between 3 and 9 dpi

In order to determine whether the presence and activity of regenerated neurons is correlated with functional recovery from SCI, we first needed to define which specific motor activity patterns are restored during the time that neurons appear in the injury site. Following SCI at 5 dpf, fictive swimming behavior was evoked by head stimulation and changes in light intensity, and two extracellular electrodes were used to monitor motor activity at 3 and 9 dpi (Fig. 1A-B). Recording electrodes placed at segments rostral to the injury informed normal activity patterns, while electrodes at injured segments and segments caudal to the injury were used to monitor recovery. Evoked swim bouts comprised multiple bursts of motor activity that were analyzed for burst duration and rostro-caudal propagation latency between ipsilateral segments (Fig. 1A), and left-right alternation latency between contralateral segments (Fig. 1B). Injured segments and caudal segments were then compared to rostral segments at 3 dpi and 9 dpi to determine if motor patterns had recovered at those timepoints.

**Figure 1.**
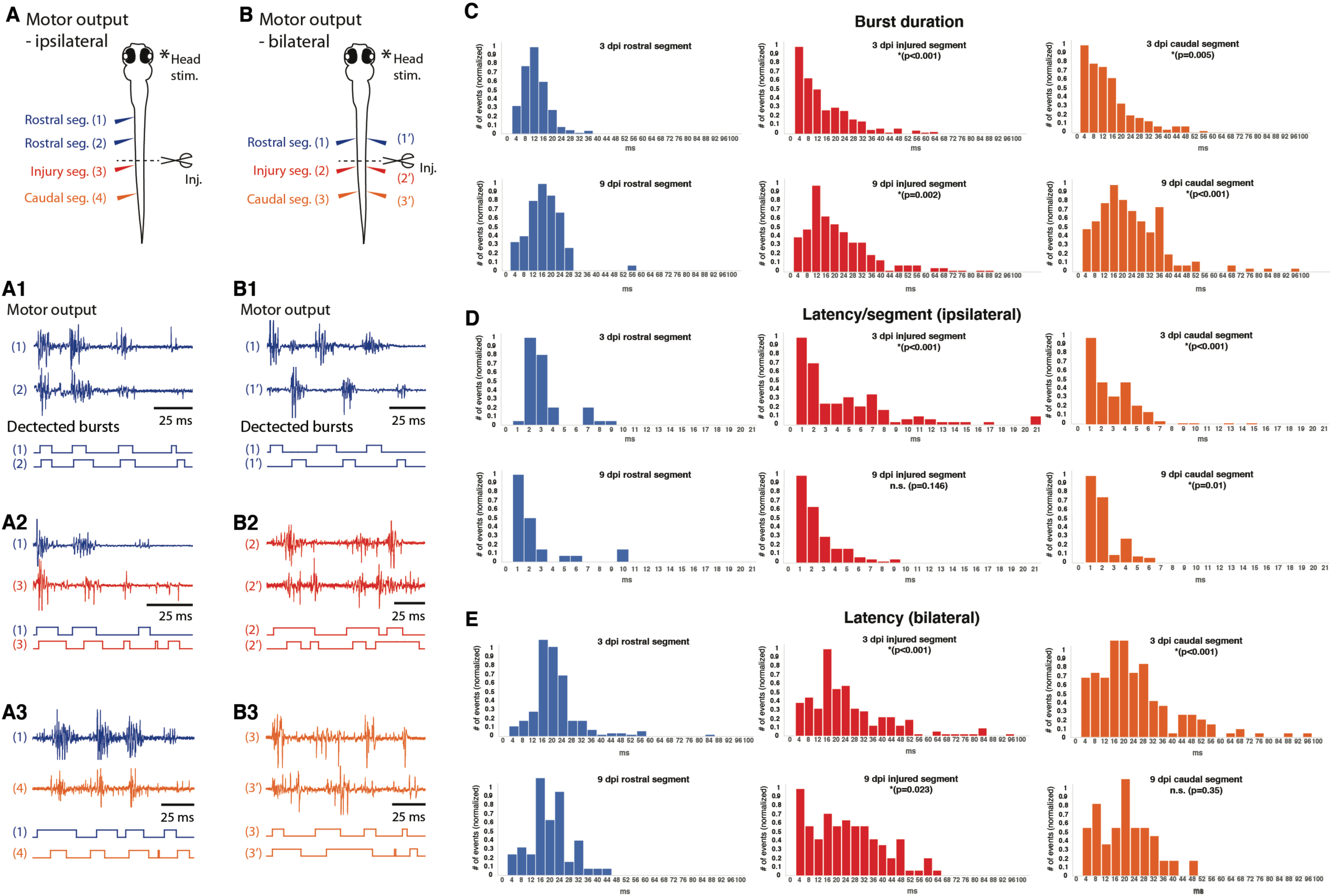
Specific locomotor patterns recover between 3-9 dpi. (A) Schematic representation of the experimental preparation for ipsilateral motor output recordings. One extracellular electrode was placed at a segment rostral to the injury [position (1)] and a second electrode was moved caudally along the body of the fish to record motor output from positions (2) to (4). (A1-3) Example traces at 9 dpi of the motor output and detected bursts at each position. Recordings from segments rostral to the injury (A1) were used to compare with recordings from injury segments (A2) and segments caudal to the injury (A3). For each panel, the top portion shows example traces of recorded motor output, while the bottom portion shows bursts detected over a set threshold, used to determine burst durations and latencies. (B) Schematic representation of preparation for contralateral motor output recordings. Extracellular electrodes were placed on both sides of the fish at the same segment rostral to the injury segment [positions (1) and (1’)], then both subsequently moved caudally to the injury segment [positions (2) and (2’)] and then to a segment caudal to the injury [positions (3) and (3’)]. (B1-3) Shows example traces at 9 dpi of the motor output and detected bursts at each position. (C) Histograms of burst durations from three ipsilateral positions in 3 dpi and 9 dpi fish. (D) Histograms of onset latencies of motor bursts between ipsilateral segments in 3 dpi and 9 dpi fish. Latency data were normalized according to the number of segments separating the recording electrodes. (E) Histograms of onset latencies of motor bursts on opposite sides of the body at the same segment in 3 dpi and 9 dpi fish. For ipsilateral recordings at 3 dpi: n=41 swim bouts; at 9 dpi: n=65 swim bouts. For bilateral recordings at 3 dpi: n=107 swim bouts; at 9 dpi: n=74 swim bouts.

At 3 dpi, the distribution of activity recorded from injured segments and caudal segments was significantly different from those recorded from rostral segments, due to more events with long burst duration and latency (Fig. 1C-E). However, by 9 dpi rostro-caudal propagation latency at injured segments was not significantly different from rostral segments (p=0.146, Fig. 1D). In addition, left-right alternation latency in segments caudal to the injury was not statistically different from rostral segments (p=0.35, Fig. 1E). While rostro-caudal propagation latency at injured segments (p=0.01) and left-right alternation latency at caudal segments (p=0.023) were still significantly different from rostral segments at 9 dpi, both measurements showed a decrease in long-latency events compared to 3 dpi (Fig. 1D, E), suggesting partial recovery in these regions. Together, our observations that specific motor activity patterns critical for rhythmic swimming behavior recover between 3 and 9 dpi is consistent with a gradual restoration of spinal cord locomotor circuitry during this time period.

We next examined the behavioral effects of recovery from SCI by analyzing tail bend wave propagation during swimming. The optomotor response (OMR) was used to evoke forward swimming behaviors in head-embedded larvae (schematic shown in Fig.2A). During repeated OMR trials we used automated behavior tracking to quantify the frequencies present in propagating swim waves both rostral and caudal to the site of injury at 3 and 9 dpi. We found that PSDs of swim waves in the physiologically relevant frequency range between 10 and 30 Hz were significantly different between uninjured and injured fish at 3 dpi, both rostral and caudal to the injury site (Fig. 2B and S1). Significant PSD differences remained in rostral regions at 9 dpi (Supplemental movies S1 and S2), consistent with studies showing a compensatory effect in neural plasticity following injury (Behrman et al 2006, Martinez et al 2011, Nishimura et al 2007). In contrast, PSDs caudal to the injury at 9 dpi were statistically indistinguishable from those in uninjured fish (Fig. 2B) suggesting a recovery of normal swim wave propagation in this region. Taken together, these data demonstrate partial recovery of locomotor behavior correlated with the reappearance of neurons that we previously observed following injury (Briona & Dorsky 2014).

**Figure 2:**
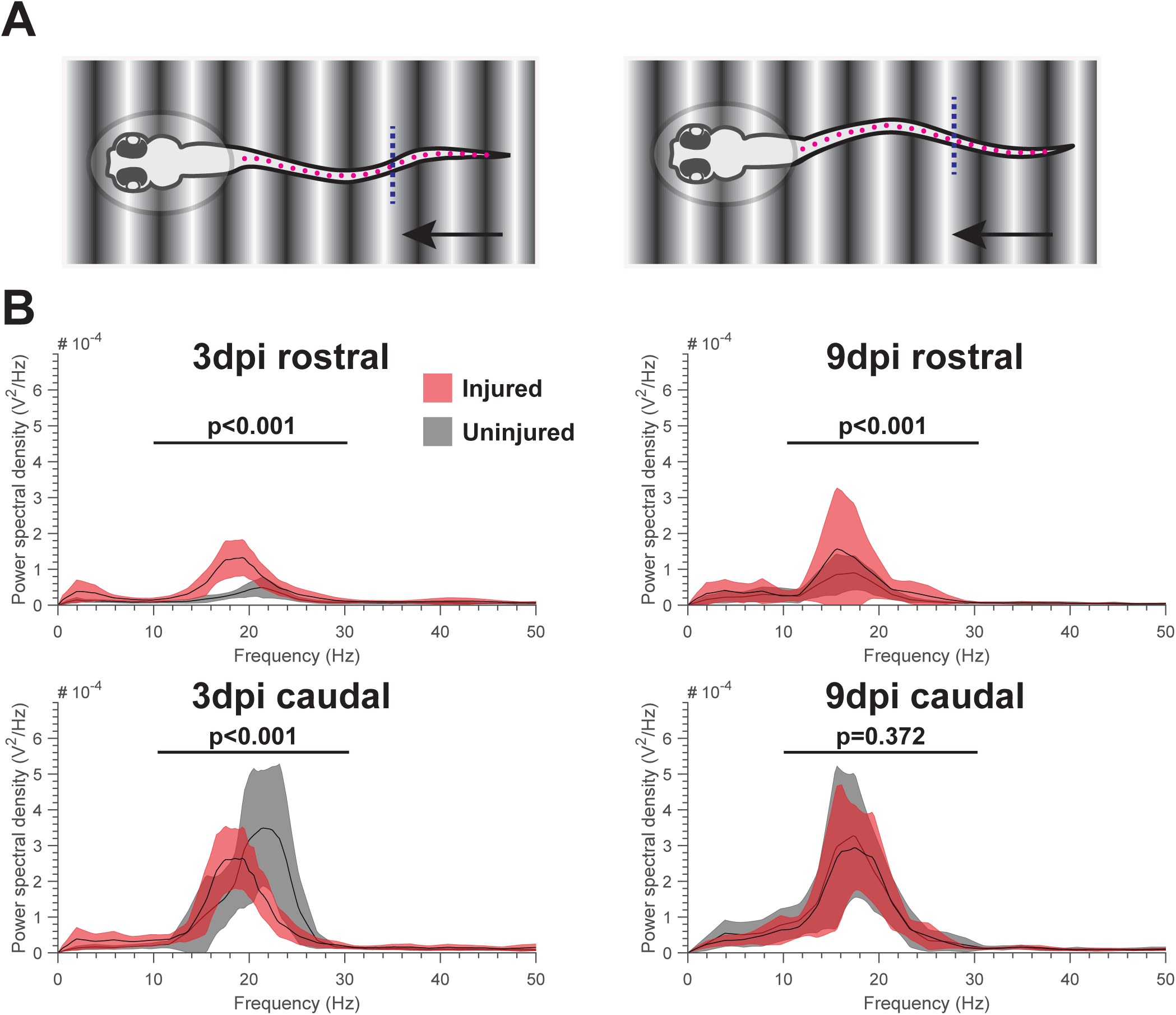
Kinematics of swimming behavior partially recovers between 3 and 9 dpi. (A) Schematic representation of the OMR assay. Moving gradient bars are projected onto the stage beneath head-embedded fish. Pink dots represent the 20 points that are used to measure tail bend angles, starting caudal to the swim bladder and ending at the tip of the tail. Blue line indicates the injury plane. Power spectral densities (PSDs) were calculated for regions rostral and caudal to the injury. (B) Power (Y-axis) is plotted against swim wave frequency (X-axis) for rostral and caudal regions of injured and uninjured fish at 3 and 9 dpi. Black lines represent mean PSDs and filled areas represent a 95% confidence interval for injured (red) or uninjured (grey) fish respectively. T-tests were performed for power at frequencies between 10 and 30 Hz comparing injured and uninjured fish. (3dpi uninjured n=16; 3dpi injured n=22; 9dpi uninjured n=20; 9dpi injured n=11)

### Newly-generated neurons repopulate the injury site between 3 and 9 dpi

To assess the potential contribution of neuronal regeneration to behavioral recovery, we next determined what proportion of neurons in the injured segment were generated after transection. Because incomplete lineage labeling in our previous analyses (Briona & Dorsky 2014, Briona et al 2015) did not allow us to answer this question, we used a pan-neuronal transgene expressing the photoconvertible fluorophore Kaede (Mutoh et al 2006) to mark all neurons that were present before injury. At 4 dpf approximately 20 spinal cord segments surrounding the prospective injury site were photoconverted from green to red (Fig. 3A-F). After SCI at 5 dpf, larvae were examined for Kaede-expressing neurons at three timepoints during the recovery period. At 1 and 3 dpi, the injured segment was completely devoid of newly-generated (green only) neurons, with only a few photoconverted (red and green) neurons and axons present (Fig. 3G-L). However, at 9 dpi, most neurons in the injured segment were green, but not red (Fig. 3 M-O), indicating that they were generated after photoconversion. In contrast, all neurons in uninjured fish still expressed red photoconverted Kaede (Fig. 3 P-R). Focusing on a defined 120*µ*m regenerative zone extending 60*µ*m rostral and caudal to the plane of injury, we found that 82% (+/-2%, S.E.M.; N=14) of neurons in this zone expressed only unconverted (green, but not red) Kaede. These data indicate that by the time of functional recovery at 9 dpi, a large majority of neurons in the injured spinal cord segment were generated after SCI.

**Figure 3.**
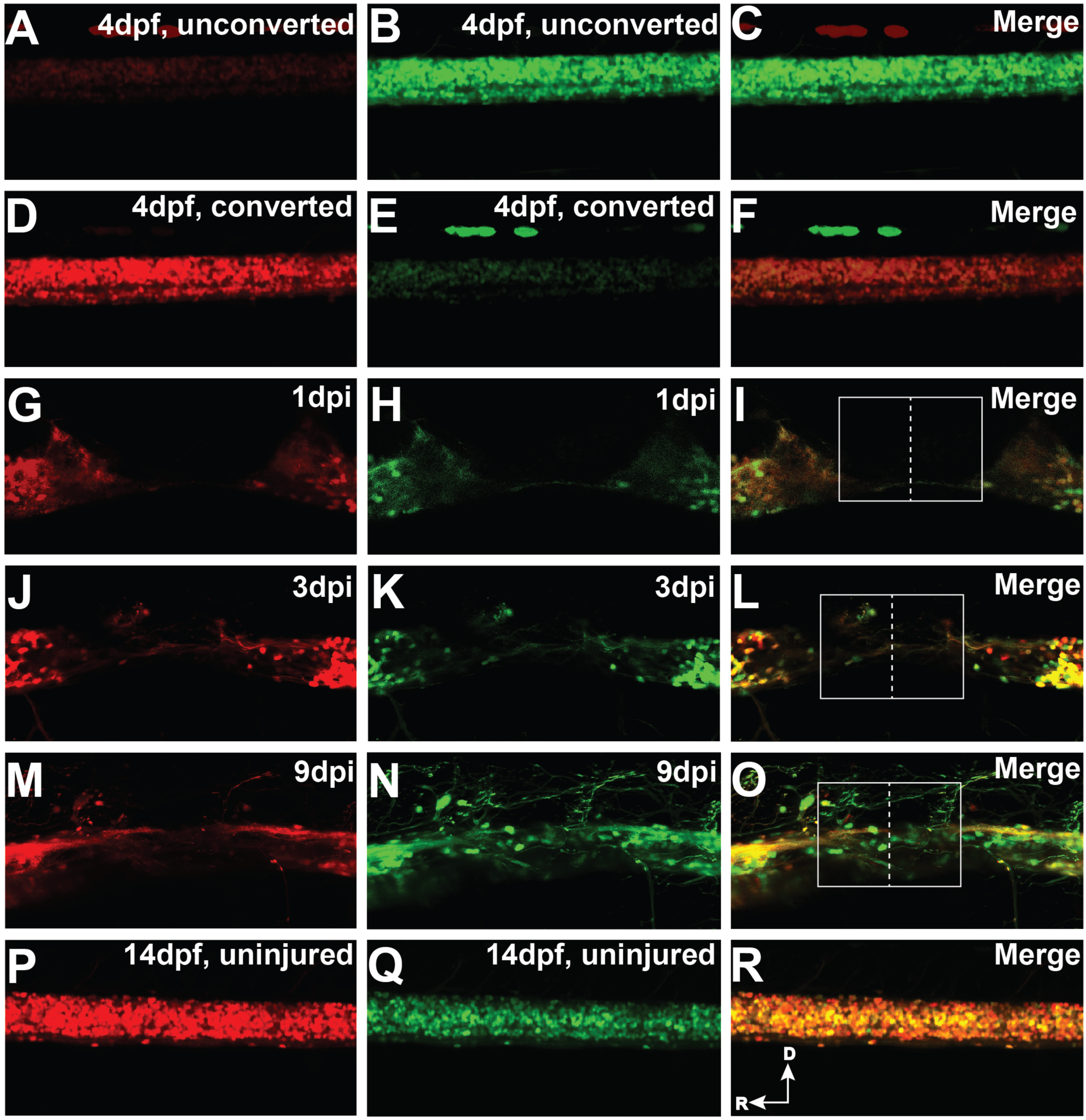
The majority of neurons repopulating transected spinal cord segments are generated after injury. (A-F) Lateral views of the spinal cord at 4 dpf showing neuronal Kaede expression before (A-C) and after (D-F) photoconversion. (G-I) Transected spinal cord at 1 dpi, showing loss of all Kaede-expressing neurons in the injured segment. (J-L) Transected spinal cord at 3 dpi showing photoconverted (red and green) Kaede-expressing axons and neurons entering the injured segment. (M-O) Transected spinal cord at 9 dpi showing photoconverted (red and green) Kaede-expressing axons and non-photoconverted (green only) neurons in the injured segment. Dashed line represents plane of injury and box represents a 120μm-wide regeneration zone. (P-R) Untransected spinal cord at 14 dpf showing perdurance of photoconverted (red and green) Kaede-expressing neurons at 14 dpf. Images are maximum-intensity projections of 100*µ*m confocal stacks.

### Interneuron subtypes involved in distinct features of swimming behavior appear at the injury site between 3 and 9 dpi

Locomotor activity requires several distinct subtypes of spinal interneurons that are conserved throughout diverse vertebrate species. These include V0 commissural interneurons that function in left-right alternation (Björnfors & El Manira 2016), V2a interneurons that provide both premotor excitation (Kimura et al 2013, McLean et al 2007, Menelaou et al 2014), and contralateral inhibition (Menelaou & McLean 2019) and V2b interneurons that provide ipsilateral inhibition for speed control (Callahan et al 2019). We therefore hypothesized that these interneuron subtypes would be regenerated during the recovery of swimming behavior between 3 and 9 dpi.

Each interneuron subtype can be identified using molecular and genetic markers in the larval zebrafish spinal cord. We labeled V0 interneurons [commissural secondary ascending (CoSA) in zebrafish] with an anti-Pax2.1 antibody (Batista & Lewis 2008), V2a interneurons [circumferential descending (CiD) in zebrafish] with a *vsx2:gal4* transgene (Kimura et al 2008) and V2b interneurons [ventral longitudinal descending (VeLD) in zebrafish] with a *gata2:gfp* transgene (Meng et al 1997). In injured segments between 3 and 9 dpi we observed the appearance of multiple Pax2.1-immunopositive nuclei (Fig. 4A-C), and GFP+ neurons in transgenic *vsx2:gal4;uas:gfp* (Fig. 4D-F) and *gata2:gfp* (Fig. 4G-I) larvae. From these observations we conclude that the interneuron subtypes necessary for modulating swim behavior appear to be produced in injured segments during the time of behavioral recovery.

**Figure 4.**
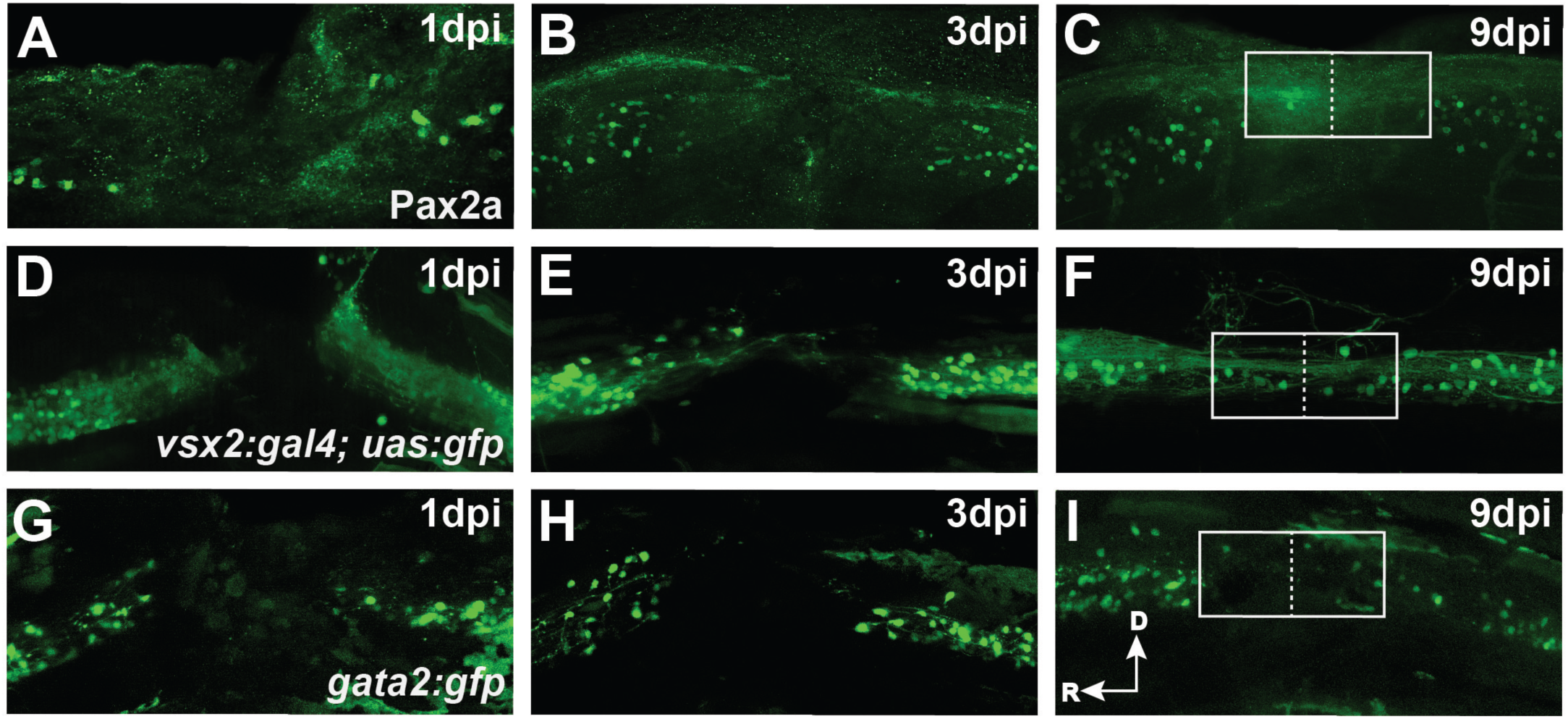
Molecularly and genetically-defined interneuron subtypes repopulate injured spinal cord segments by 9 dpi. (A-C) Immunohistochemical labeling shows the presence of Pax2a+ V0 interneurons in injured segments by 9 dpi. (D-F) GFP expression in *vsx2:gal4;uas:gfp*+ larvae shows the presence of V2a interneurons in injured segments by 9 dpi. (G-I) GFP expression in *gata2:gfp*+ larvae shows the presence of V2b interneurons in injured segments by 9 dpi. Dashed line represents plane of injury and box represents a 120μm-wide regeneration zone. Images are maximum-intensity projections of 100*µ*m confocal stacks.

### *vsx2:gal4+ i*nterneurons in injured segments are physiologically active at 9 dpi

The reappearance of defined interneuron subtypes by 9 dpi led us to ask whether they are capable of participating in functional circuitry at that timepoint. We focused our analysis on *vsx2+* V2a interneurons because their physiological properties and role in swim behavior have been extensively characterized in larval zebrafish (Ampatzis et al 2014, Eklof-Ljunggren et al 2012, Menelaou & McLean 2019, Menelaou et al 2014, Song et al 2018). To visualize their full morphology, fluorescent dye was used to fill single GFP+ cells in uninjured (n=18) and injured (n=11) *vsx2:gal4;uas:gfp* larvae (Fig. 5A-B). All GFP+ neurons in uninjured 14 dpf larvae exhibited typical V2a morphologies previously documented in zebrafish (Kimura et al 2008, Menelaou et al 2014) with axons that projected ventrally, then caudally across multiple segments and lateral dendritic processes at the level of the cell body (Fig. 5A). In contrast, GFP*+* neurons found in injured segments at 9 dpi had highly variable morphologies with processes extending rostrally and caudally (Fig. 5B), while stereotypic ventro-caudal projecting axons were not observed. In addition, the soma size of GFP+ neurons in injured segments was significantly smaller than cells in the equivalent segments of uninjured larvae (Fig. 5C).

**Fig 5.**
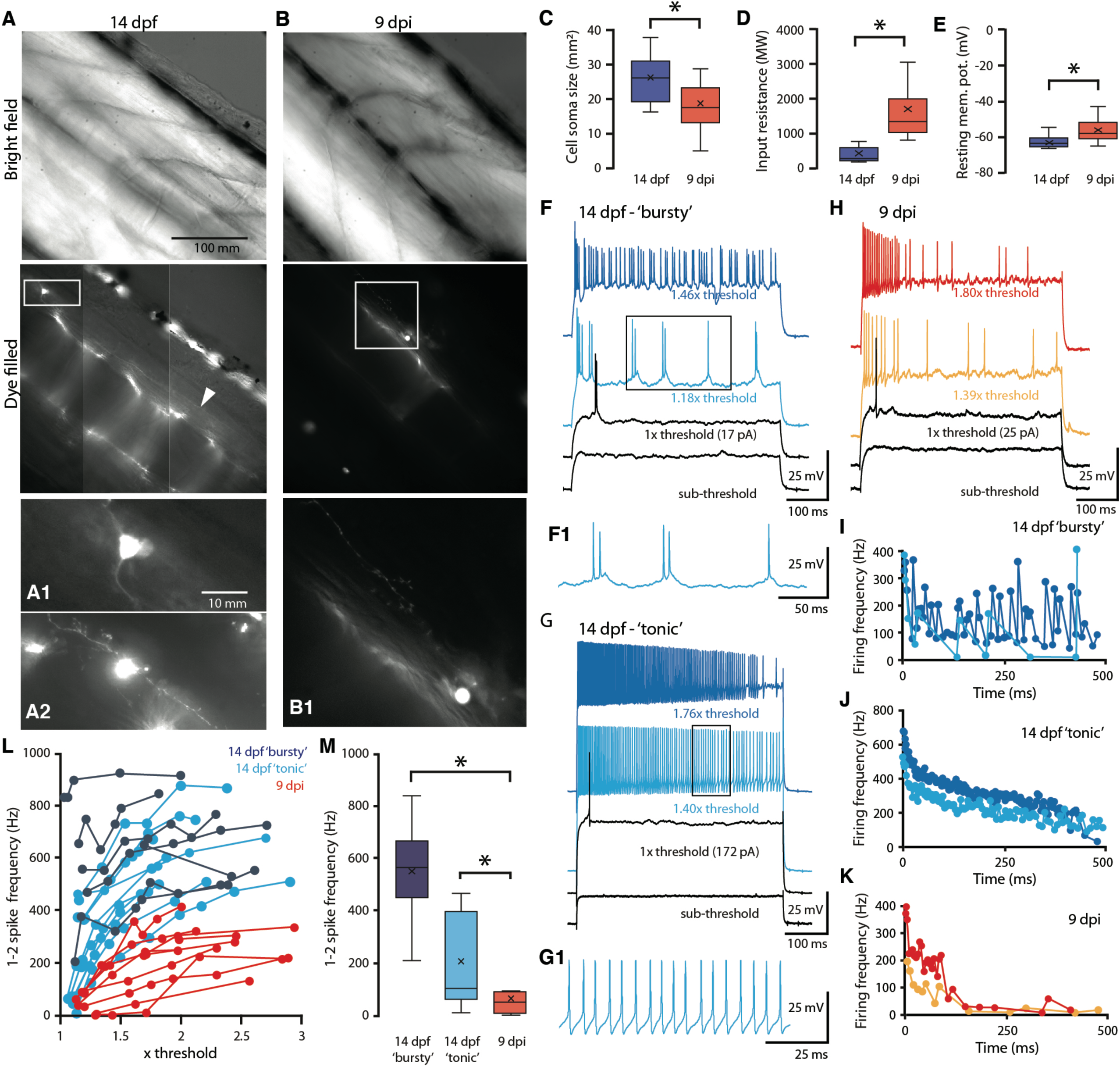
*vsx2:gal4+* interneurons in regenerated segments produce action potentials at 9 dpi. (A-B) Images of a GFP+ neuron in uninjured *vsx2:gal4;uas:gfp* larvae at 14 dpf (A) and a GFP+ neuron in a regenerated segment at 9 dpi (B). Top panels show brightfield images revealing distinct chevron-shaped muscle segments, middle panels show the same field of view with florescence imaging in which a single GFP+ neuron has been filled with dye. White boxes in middle panels have been enlarged in bottom panels. Uninjured GFP+ neurons project axons ventrally, then caudally (A1) across multiple segments (arrowhead in middle panel) and often have dendritic processes at the same level of the cell body (A2). At 9 dpi, GFP+ neurons in regenerated segments have processes that are highly variable in projection pattern and lack the characteristic ventral-caudal projecting axons found in uninjured cells (B1). GFP+ neurons in regenerated segments are smaller (C), have a higher input resistance (D) and are more depolarized at rest (E) compared to uninjured cells. (F) Example traces of a ‘bursty’ type GFP+ neuron in response to depolarizing current steps of increasing amplitude. Bursty neurons fire clusters of spikes rhythmically with lower frequencies between clusters (F1). (G) Example traces of a ‘tonic’ type GFP+ neuron in response to depolarizing current steps of increasing amplitude. Tonic neurons fire continuously with a more consistent firing frequency (G1). (H) Example traces of a GFP+ neuron from a regenerated segment in response to depolarizing current steps of increasing amplitude. (I-K) Change in firing frequencies in response to 500 ms of the depolarizing current steps at different amplitudes over threshold for the three GFP+ neurons shown in F-H. (L) 1-2 spike frequency in response to current steps at different amplitudes over threshold for all cells, showing the lower spike frequencies of GFP+ neurons in the regenerated segment for most amplitudes (red). (M) 1-2 spike frequencies in response to current steps closest to threshold, showing that GFP+ neurons in the regenerated segment have significantly lower spike frequencies compared to either type of uninjured GFP+ cells.

Whole-cell patch clamp recordings showed that GFP*+* neurons in injured segments at 9 dpi (n=24) had significantly higher input resistances (Fig. 5D) and were more depolarized at rest compared to GFP+ neurons in uninjured larvae (n=18) (Fig. 5E). Depolarizing current steps were applied to determine firing thresholds, and current steps of increasing amplitude were applied to examine spiking behavior at different magnitudes above threshold (Fig. F-H). GFP*+* neurons in uninjured larvae responded to suprathreshold current injections with either clusters of high frequency spikes (“bursty”, Fig. 5F, I), or with sustained firing that gradually decreased with time (“tonic”, Fig. 5G, J). Strikingly, we found that all GFP+ neurons in injured segments at 9 dpi responded to suprathreshold current injections with spiking behavior that decreased with time, although they could not be classified as either bursty or tonic (Fig. 5H, K). GFP+ neurons in regenerated segments had lower firing frequencies between the first and second spikes (1-2 spike frequency) for all current steps at amplitudes above threshold, and had significantly lower spike frequencies around threshold, compared to both bursty and tonic types of GFP+ neurons in uninjured larvae (Fig. 5L, M). Together these data indicate that while the morphology and physiology of neurons generated after SCI does not precisely resemble genetically comparable cells in uninjured larvae at the same stage, they are capable of responding to excitatory input with action potentials by 9 dpi.

### *vsx2:gal4* interneurons in injured segments are activated in phase with motor output at 9 dpi

If spinal interneurons generated after SCI can participate in the recovery of swimming behavior, their activity should be synchronized with motor output. This was assessed by performing whole-cell patch clamp recordings from GFP*+* neurons in *vsx2:gal4;uas:gfp* larvae along with simultaneous peripheral motor nerve recordings (Fig. 6A, B). In both injured and uninjured larvae, rhythmic depolarization of the membrane potential in GFP+ neurons corresponded with swim bursts (Fig. 6C, D). GFP+ neurons in uninjured larvae reproducibly fired during a high percentage of swim bursts per swim bout (n=33 bouts, Fig. 6E). In contrast, only some GFP+ neurons in injured segments fired during most swim bursts, while others fired during very few (n=44 bouts, Fig. 6E). Overall, GFP+ neurons in uninjured larvae had a higher firing probability during swim bouts (Fig. 6F), and generally fired more spikes per burst (Fig. 6G) compared to GFP+ neurons in injured segments. Despite these physiological differences, GFP+ neurons in injured segments showed clear synchronization with motor activity by 9 dpi.

**Fig 6.**
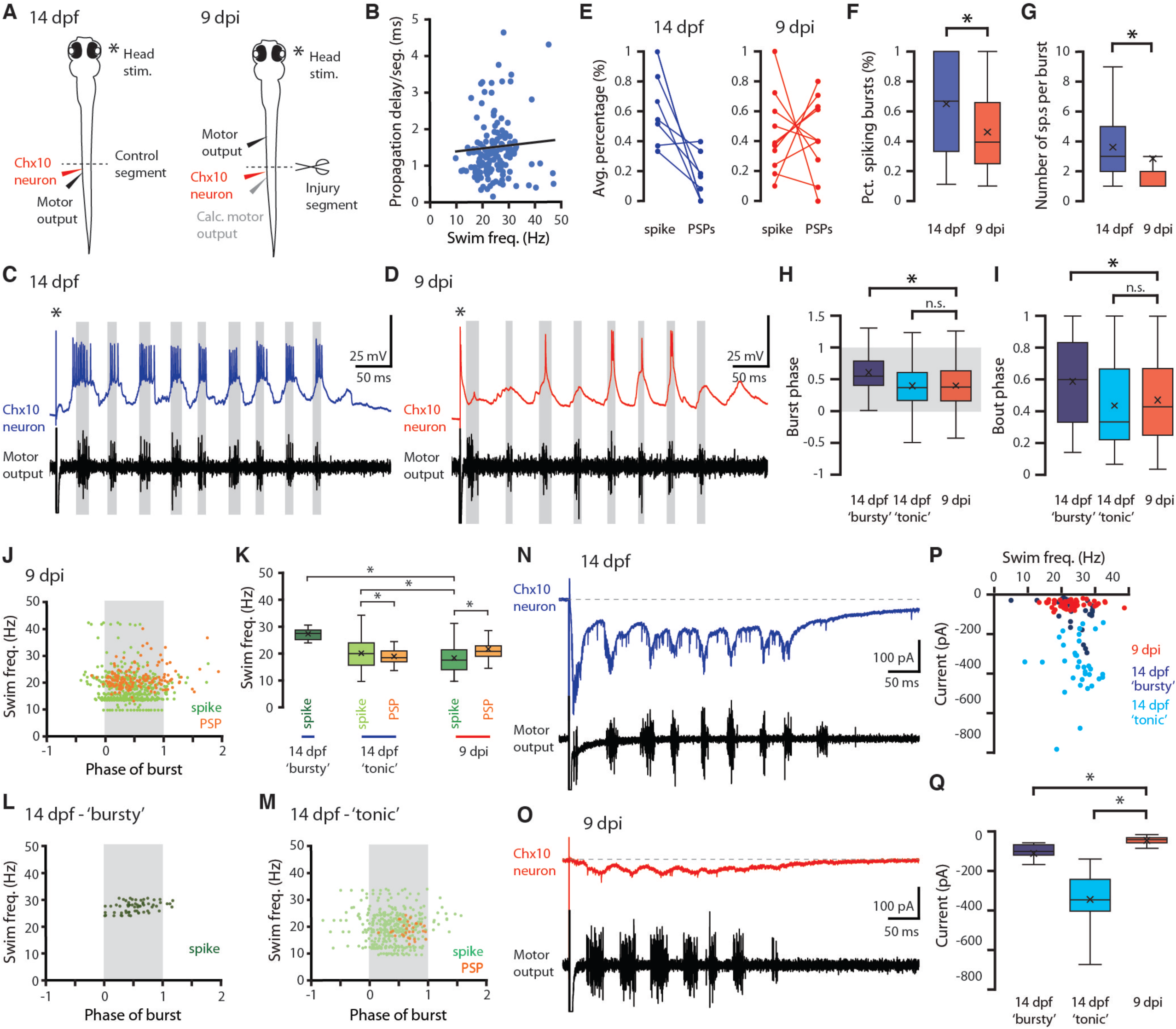
*vsx2:gal4+* interneurons in regenerated segments receive excitatory input and are activated in phase with motor activity. (A) Schematic representation of the experimental preparation for simultaneous patch-clamp recordings of GFP+ neurons and motor output recordings from peripheral motor nerves in *vsx2:gal4;uas:gfp*+ larvae. In uninjured fish, motor output was recorded from the same segment as neuron recordings, while in injured fish motor output was recorded from segments rostral to the injury in order to obtain motor patterns that had not been compromised by injury. (B) The propagation delay per segment was calculated with 14 dpf uninjured fish (1.48 ± 1.18 ms/segment, n=26 swim bouts) and used to derive the motor output at the injury segment from the recorded motor output at segments rostral to the injury in injured fish. (C,D) Example traces of GFP+ neurons and motor output in response to a brief electrical stimulus. Asterisk denotes the stimulus artifact and gray boxes demark the duration of each motor burst. (E) Average percentage of motor bursts associated with either spikes or depolarizations in uninjured and injured fish. (F) Firing probability for GFP+ neurons during a swim bout. (G) Spikes per burst fired by GFP+ neurons in injured segments. (H,I) Phase of GFP+ neuron firing during motor bursts and bouts. Gray box in (H) denotes the beginning and end of a motor burst. (J) Distribution of GFP+ neuron spike and depolarization timing at different swim speeds in injured fish. Gray bars denote the beginning and end of a motor burst. (K) Comparison of swim frequencies at which GFP+ neurons were activated. (L, M) Distribution of GFP+ neuron spike and depolarization timing at different swim speeds for ‘bursty’ and ‘tonic’ types in uninjured fish. Gray bars denote the beginning and end of a motor burst. (N, O) Example traces of excitatory inward current received during a swim bout for GFP neurons. (P, Q) Maximum (P) and average (Q) amplitude of inward current received per swim bout for GFP+ neurons.

When we analyzed burst and bout phase, defined as the timing of a spike within a single burst or swim bout, respectively, we found that GFP+ neurons in injured segments fired at the same phase of bursts and bouts as tonically-firing neurons in uninjured larvae. This was in contrast to bursty neurons, which fired later in bursts and bouts (Fig. 6H, I). Interestingly GFP+ neurons in injured segments fired at lower swim frequencies, whereas higher swim frequencies more often corresponded to depolarizations but not spiking activity (Fig. 6J, K). In uninjured larvae tonically-firing GFP+ neurons fired across swim frequencies, while bursty neurons fired at higher swim frequencies (Fig. 6K-M).

Excitatory drive during swimming measured through voltage-clamp recordings demonstrated that GFP+ cells in both injured segments and uninjured larvae received rhythmic excitatory inputs that corresponded with swim bursts (Fig. 6N, O). However, GFP+ neurons in injured segments received significantly less excitatory drive compared to GFP+ neurons in uninjured larvae, regardless of swim frequency (Fig. 6P, Q). Despite these differences, the fact that interneurons generated following SCI are activated in phase with evoked motor output by 9 dpi suggests that they are becoming integrated into locomotor circuitry.

## Discussion

### Regeneration of defined interneuron subtypes is correlated with partial behavioral recovery

In this study we show for the first time that newly-generated interneurons are physiologically active during functional recovery from SCI. Following spinal cord transection of zebrafish larvae, newly-generated V0, V2a, and V2b interneurons, which are critical for various aspects of routine swim behaviors, repopulate the injured region between 3 and 9 dpi. The function of these interneuron subtypes in zebrafish and other vertebrates indicates that they play important roles in the central pattern generator network of the spinal cord (Berkowitz et al 2010, Eklöf-Ljunggren et al 2012, Fetcho et al 2008), which regulates rhythmic left-right alternations of motor activity necessary for forward swimming (Wiggin et al 2012, Wyart et al 2009). Specific aspects of swimming behavior also begin to recover during the same time period, including the latency of wave propagation at the injured segment, and left-right alternation and swim wave frequencies caudal to the injured segment. While the propagation of motor activity through an injured spinal cord segment could theoretically be accomplished by axon regrowth from surviving neurons rostral and caudal to the injury site, the regenerated interneurons that we observed may also play a role in this process. Given the known roles of V0, V2a, and V2b interneurons in both ipsilateral motor propagation and contralateral inhibition (Björnfors & El Manira 2016, Callahan et al 2019, Menelaou & McLean 2019), it is reasonable to hypothesize that regeneration of these neuronal subtypes helps to refine the kinetics of swimming after injury.

### Newly-generated interneurons integrate into locomotor circuitry

In this study we show that by 9 dpi *vsx2:gal4+* interneurons in injured spinal cord segments receive excitatory inputs, are capable of producing action potentials following depolarization, and fire synchronously with evoked motor behavior. To our knowledge this is the first demonstration of endogenously-generated spinal cord neurons integrating into locomotor circuits following SCI. While the properties of these cells clearly differ from V2a interneurons in uninjured spinal cord segments, it is possible that they continue to mature beyond 9 dpi and eventually adopt characteristic bursty or tonic physiology. In addition, we do not know if or when newly-generated V2a interneurons make appropriate efferent connections with motor neurons in caudal segments, partly due to a lack of characteristic axonal projections at 9 dpi. Stimulation of these neurons paired with optical or electrophysiological recording of activity from caudal motor pools at different timepoints could help resolve this question.

### A potential role for local neurogenesis in rhythmic locomotor activity

Our behavioral and physiological analyses suggest that even though some aspects of signal propagation down the spinal cord recover by 9 dpi, other aspects of motor behavior and neuronal physiology do not fully recover by this timepoint. Although it is possible that this is a permanent physiological effect of the injury, continued refinement of connectivity and activity over time may eventually make motor patterns indistinguishable from uninjured fish. By testing the necessity of newly-generated neurons for behavioral recovery using genetic manipulations, future studies can determine whether they are essential for the recovery of normal swim behaviors.

## Acknowledgements

We thank the University of Utah Centralized Zebrafish Animal Resource for zebrafish husbandry, and the University Cell Imaging Core Facility for microscopy support. We thank Angie Serrano for assistance with confocal imaging, Kristen Kwan for providing the Pax2a antibody, and David Grunwald for helpful discussions and critical reading of the manuscript.

